# A general-purpose mechanism of visual feature association at the core of word learning

**DOI:** 10.1101/2020.07.14.202176

**Authors:** Yamil Vidal, Eva Viviani, Davide Zoccolan, Davide Crepaldi

## Abstract

Writing systems are a recent cultural invention, which makes it unlikely that specific cognitive mechanisms have developed through selective pressure for reading itself. Instead, reading might capitalize on evolutionary older mechanisms that originally supported other tasks. Accordingly, animals such as baboons can be trained to perform visual word recognition. This suggests that the visual mechanisms supporting reading might be phylogenetically old and domain-general. Here we propose that if the human reading system relies on domain-general visual mechanisms, effects that are typically found within the domain of reading should also be observable with non-orthographic visual stimuli. To test this hypothesis, we systematically tested different types of visual material with the same experimental design. Subjects were passively familiarized with a set of composite visual items, and then tested in an oddball paradigm for their ability to detect novel stimuli. Some of these novel stimuli shared their statistical structure with the familiar items, and were found to be hard to detect in two experiments using strings of letter-like symbols; this replicates the well-known, and supposedly reading-specific, bigram effect. Crucially, in two further experiments we show that the same effect emerges with made-up, 3D objects and sinusoidal gratings. The effect size was equivalent across experiments, despite the use of radically different stimuli. These data suggest that a fundamental mechanism behind visual word learning also supports the learning of other visual stimuli, implying that such mechanism is general–purpose. This mechanism would enable the statistical learning of regularities in the visual environment.

## 2 Introduction

As writing systems are a relatively novel invention (slightly over 5K years ago [1]), they could not have influenced the evolution of our species. Despite this, part of the human cerebral cortex seems to be specialized for reading. For example, during reading acquisition, cortical surface in the left fusiform gyrus that is originally devoted to the processing of visual objects (and faces in particular) becomes tuned to orthographic material [2–7].

One possible reason for this might be that cultural inventions such as writing hijack or recycle older domain-general cognitive processes [2, 8]. In line with this, it has been shown that baboons can be trained to distinguish words from nonwords based on orthographic regularities in letter cooccurrence [9], and generalize to new unseen stimuli with above chance performance. These results were later replicated with pigeons [10], which suggests that visual systems with a radically different organization compared to that of primates can support the processing of orthographic regularities. Furthermore, recent work has shown that neurons in high-level visual cortex of naive macaque monkeys could implement orthographic processing tasks [11]. This mounting body of evidence shows that part of what is usually considered reading-specific processing can be performed by domain-general visual mechanisms present in non-linguistic animals. This suggests that such visual mechanisms might be phylogenetically old and precede the emergence of written language.

Here we propose that if the human reading system indeed relies on domain-general visual mechanisms, then some of the effects that are typically found with orthographic material should be observable with non-orthographic visual stimuli. To this aim, in our study we systematically tested different types of visual material with the exact same experimental design, in search of an effect that is typically studied in the context of orthographic processing, i.e., participants’ sensitivity to bigram frequencies (frequency of occurrence of pairs of graphemes).

Bigram frequencies have been proposed to be encoded as an intermediate step between single graphemes and small words or word fragments (e.g., the word HOUSE is coded as a collection of its bigrams HO, OU, US, SE [12–14]). While the role of bigram frequencies in reading is open to debate (see [15] and [16] for two opposite views), fMRI [17, 18] and more recently human intracranial EEG recordings [19] have shown that populations of neurons in the left fusiform gyrus are sensitive to this factor. This, together with the aforementioned animal research literature which relies on the manipulation of bigram frequencies [9–11], makes sensitivity to bigram frequencies an ideal effect to be replicated with non-orthographic visual stimuli. Therefore we performed four experiments testing for this sensitivity, using visual stimuli whose features progressively departed from orthographic material.

Our task was based on a visual oddball design. Participants were presented with one visual stimulus at a time, 90% of which were frequent (i.e., standard) and 10% of which were infrequent (i.e., deviant). After an initial learning block in which only standard stimuli were presented, participants were asked to classify the stimuli as either “Correct” (standard) or ‘‘Mistaken” (deviant) by pressing one of two keys on a keyboard.

Within each experiment, all presented stimuli were composed by different combinations of the same constituent visual features. Importantly, two different types of deviants were used. The first type was composed by pairs of visual features that were also present in the standard stimuli. The second type was instead composed by pairs of visual features that never occurred in the standard stimuli. We will refer to these deviants as “high pair-frequency” and “low pair-frequency”, respectively. In this way, if participants are sensitive to the frequency with which specific pairs of visual features appear together, high pair-frequency deviants should be harder to detect than low pair-frequency deviants.

In experiments 1 and 2, stimuli were novel strings of pseudofonts; made-up symbols that resemble real letters in their basic visual features (e.g., number and type of strokes, number of junctions and terminations; 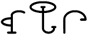). This allowed us to test whether our design was able to capture participants’ sensitivity to mean bigram frequency with orthographic-like stimuli. The use of pseudofont guarantees that results only depend on visual/orthographic processing, as artificial scripts lack any connection with phonology or meaning. Furthermore the fact that participants were completely unfamiliar with the experimental material, allowed us to avoid the influence that their particular history of exposure to orthographic material would have had in the results [e.g., 15, 20-23].

Instead, stimuli in experiment 3 consisted of images of Y-shaped visual objects with distinctive shapes attached to their terminals. These objects differed from written words in two ways. First, rather than being formed by adjacent but independent graphemes, the parts conforming the objects were physically connected in a unit. Second, the constituent parts followed a radial spatial arrangement, rather than the linear spatial arrangement that is typical of orthographic material. In experiment 4, we used even more abstract and less word-like visual stimuli – circular sinusoidal gratings in which the defining features were spatial frequency, orientation and contrast. These stimuli further differed from orthographic material, as they were devoid of any spatial arrangement and were composed solely by low level visual features. Details about the stimuli and examples can be found in figures 1 and 2 of the Results section.

**Figure 1:**
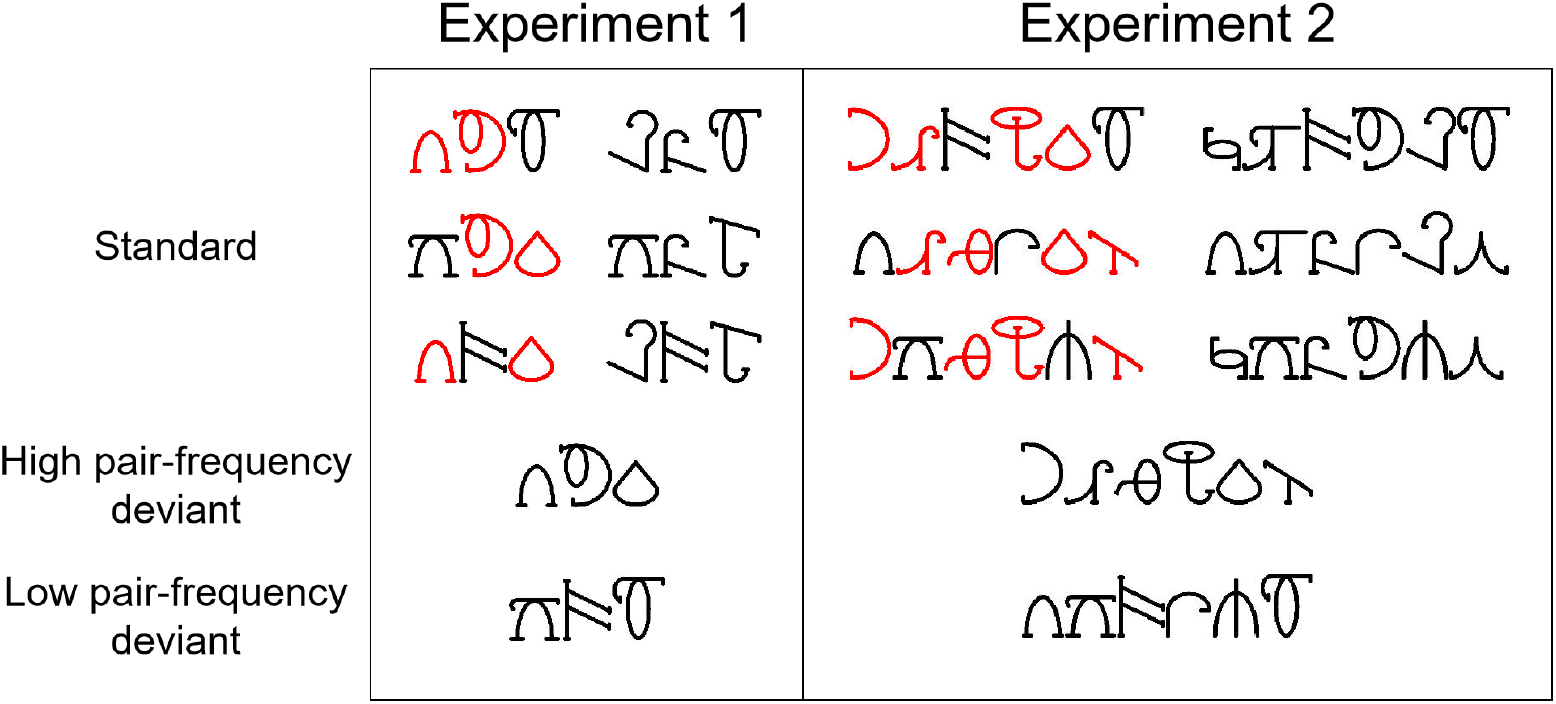
Representative stimulus sets used in experiments 1 and 2. Note that all the characters pairs making up high pair-frequency deviants are shared with standard words (marked in red for illustration purposes).

**Figure 2:**
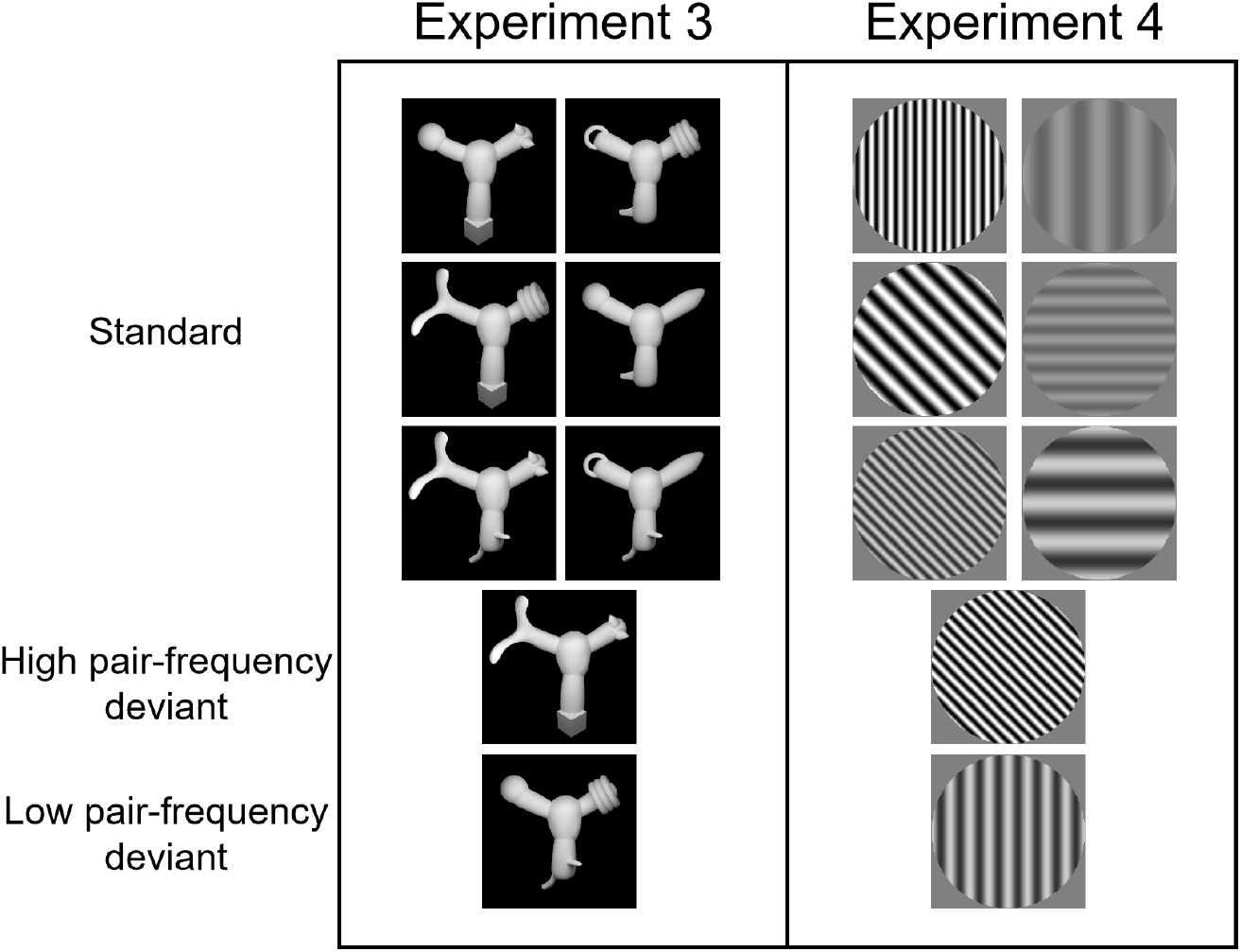
Representative stimulus sets used in experiments 3 and 4. In the case of experiment 3, each pair of shapes composing high pair-frequency deviant objects was shared with a standard object. Whereas in experiment 4, each pair of visual feature values defining high pair-frequency deviant gratings was shared with a standard grating.

In all these experiments, the same pattern of results emerged. Participants systematically found harder to reject high pair-frequency deviants, therefore showing a sensitivity to the frequency of cooccurrence of features defining the stimuli. Importantly, the size of this effect was equivalent across experiments, despite the fact that the stimuli were radically different.

## 3 Results

### 3.1 Participants are sensitive to the bigram frequencies of orthographic-like stimuli

As we anticipated in the Introduction, experiments 1 and 2 used words written using pseudofont as stimuli. That is, symbols that do not belong to the script that participants would read every day, but are similar enough to real letters and thus lend themselves to be interpreted as a novel script. We used the Brussels Artificial Character Set (BACS [22], see Figure 1 for examples).

In experiment 1, a different subset of 9 BACS characters was randomly selected to construct the stimuli for each participant. These characters were used to build 6 three-character combinations (e.g., 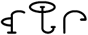), which were used as *standard* strings (they can be conceived as words in a novel lexicon that the participants had to learn). Next, taking these standard words as a base, two different *deviant* words were constructed. A high pair-frequency deviant was constructed using bigrams (pairs of characters) that were all present in the standard words. For example, the deviant 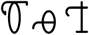 is made up of the first bigram from the standard word 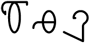, the second bigram from the standard word 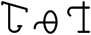 and the open bigram [12] from the standard word 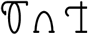. A low pair-frequency deviant was instead constructed using bigrams that were never present in standard words. For example, the deviant 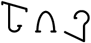 is made of characters present in the aforementioned standard words but in a unique combination. In brief, all words were constructed using the same individual characters, but while standard words and high pair-frequency deviants shared bigrams, low pair-frequency deviants did not. The same procedure was used to construct the stimuli in experiment 2, with the difference that 18 characters were randomly picked for each participant and that words were six-character long. An exemplar stimulus set for each experiment can be found in Figure 1.

Participants first completed a learning block in which only standard stimuli were presented. During this block they were instructed to pay attention and try to learn the stimuli. Stimuli were presented one at a time and remained on screen for 1.5 to 2 seconds. No explicit discrimination task was administered - the stimuli were simply viewed passively. After the learning block, the experiment followed a visual oddball design. Participants completed 6 testing blocks, where standard stimuli were presented intermixed with deviant stimuli. In these blocks, participants were given a maximum of 2 seconds to classify each stimulus as either “Correct” (standard, i.e., seen during the learning block) or “Mistaken” (deviant, i.e., not seen during the learning block) by pressing one of two keys on a keyboard. Participants were not informed about the existence of two different types of deviants and about the amount of deviants that would be presented. In these testing blocks, standard words were presented in 90% of the trials (15% each of the six items) and deviants were presented in 5% of the trials each.

The use of an oddball design, together with the way in which the stimuli were constructed, allowed us to separately manipulate two independent variables. On the one hand, the frequency of occurrence of each individual word (token frequency) was 15% for the standard stimuli and 5% for each deviant stimuli. This is the basis on which participants could carry out the task - they had to detect the low-frequency, deviant stimuli (i.e. the oddballs). On the other hand, the mean pair-frequency of each word was high for standard stimuli (5.27%) and high pair-frequency deviants (6.66%), but low for low pair-frequency deviants (1.66%). This is due to the fact that while high pair-frequency deviants shared all their bigrams with the standard stimuli, the pairs of features composing low pair-frequency deviants were only presented when such deviants were presented. In experiment 2, words were composed of 6 characters, thus all pair frequencies were exactly 1/3 of the frequencies in experiment 1. Critically, however, the ratio of mean pair-frequencies across deviants remained equal. The statistical structure of the stimuli sets was based on the one used in [24], where the authors show how the frequency of pairs of syllables affects word recognition in the context of speech segmentation.

Our task requested participants to distinguish words by their token frequency (15% for standards and 5% for deviants); therefore, high and low pair-frequency deviants should be equally rejected. However, if participants are sensitive to the frequency with which pairs of characters appear together, the detection of deviants that share bigrams with the standard words (i.e., the high pair-frequency deviants) should be harder than the detection of deviants composed by unique character combinations (i.e., the low pair-frequency deviants).

We characterized participants’ performance by computing their Sensitivity Index or d-prime score (*d*’). This is a Signal Detection Theory statistic that takes into account possible response biases [25]. All effect sizes reported are Hedges’ *g* [26, 27]. Confidence intervals reported between square brackets are 95% CI.

While the *d*’ for the high pair-frequency deviant was on average 0.84 [0.24, 1.45], it was 2.02 [1.44, 2.59] for the low pair-frequency deviant. Therefore, high pair-frequency deviants were harder to detect. The difference in *d’* between deviants was 1.17 [0.60, 1.74] (*t*_(21)_ = 4.30, *p* = 0.00016, *g* = 0.86 [0.36, 1.37]). Interestingly, as it can be seen in Figure 3A, this effect was present in the majority of the participants (86.36% [65.09%, 97.09%], or 19 out of 22, one-sided binomial test: *p* = 0.00043), which implies that the effect is highly reliable.

**Figure 3:**
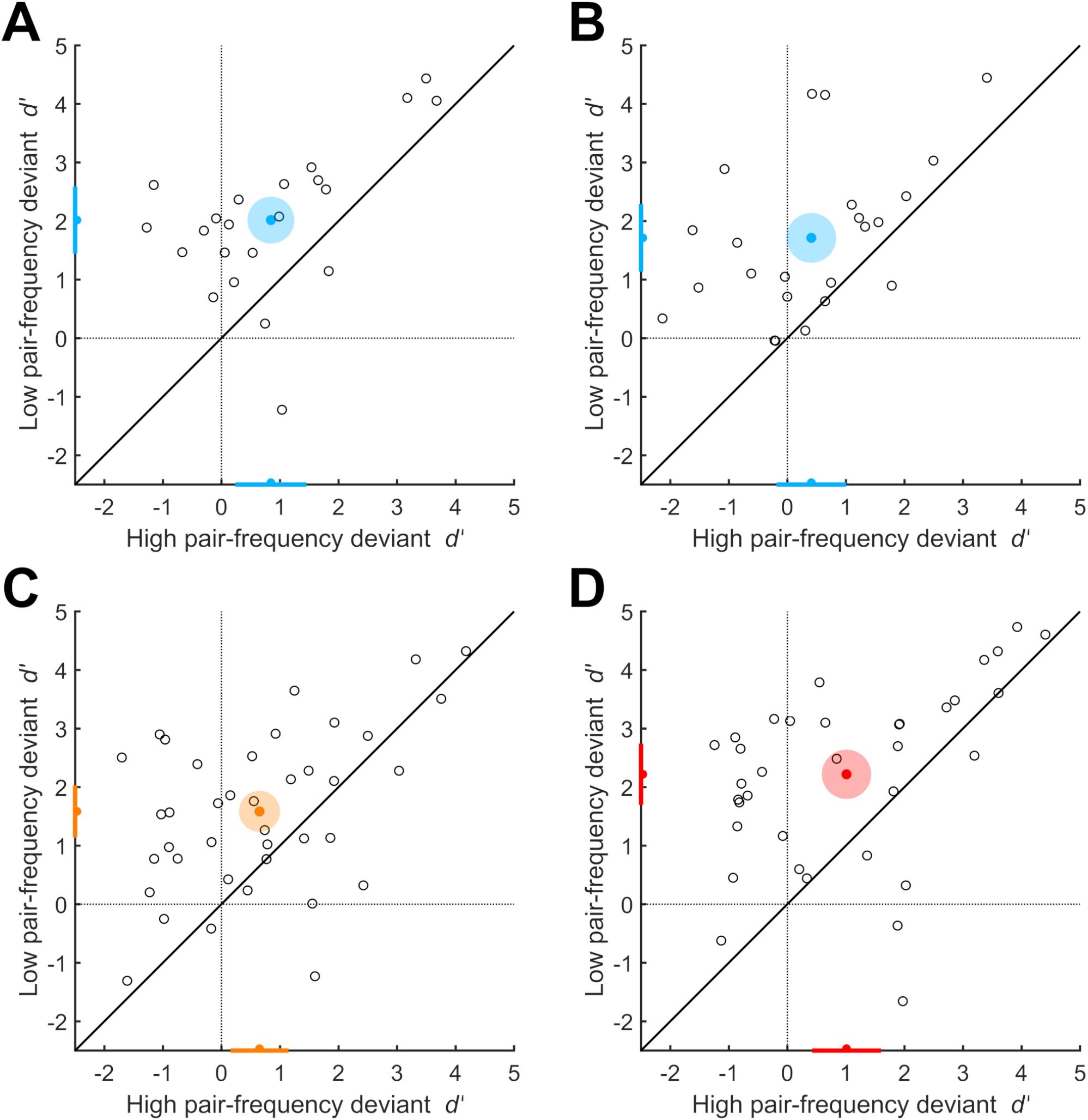
Participants are sensitive to the co-occurrence of visual features across different types of stimuli. Scatter plots of participants’ sensitivity indexes (*d*’). On each graph, the *x* and *y* axes represent sensitivity (*d*’) to high pair-frequency and low pair-frequency deviants respectively. While each dot represents a participant, the coloured dot represents the mean performance of the group. The shaded area around the mean performance denotes the group-level within participants 95% CI. Projected on each axis, a coloured dot indicates the mean performance for the respective deviant, while the error bar represents a 95% CI. Note that the majority of the participants in all of the experiments lay above the diagonal, indicating that deviants which contained pairs of features that were also present in the standards were consistently harder to detect. A: Experiment 1 (3 character word-like stimuli). B: Experiment 2 (6 character word-like stimuli). C: Experiment 3 (visual objects). D: Experiment 4 (sinusoidal gratings). A summary of participants’ hit and false alarm rates in all experiments can be found in Table S1.

The stimuli used in experiment 1 were three-character long, which implies both advantages and disadvantages. One one hand, given the short length of the stimuli, the fact that participants were sensitive to the frequency with which pairs of characters appear together, rather than encoding for whole words as a unit, is even more surprising. On the other hand, three-character words are not particularly representative of the typical length of words in real languages. To overcome this limitation, the novel words used in experiment 2 were six-character long.

Once more, the detection of high pair-frequency deviants turned out to be more challenging. Average *d*’ were 0.41 [-0.19, 1.00] for the high pair-frequency deviant and 1.71 [1.13, 2.29] for the low pair-frequency deviant. The difference in *d*’ between deviant types was 1.30 [0.70, 1.91] (*t*_(22)_ = 4.49, *p* = 9.1e-05, *g* = 0.94 [0.41, 1.48]). This effect was present in 86.96% [66.41%, 97.22%] of the participants (20 out of 23, one-sided binomial test: *p* = 0.00024). See Figure 3B.

In both experiments, it was harder to detect deviants that shared pairs of characters with the standard words. This implies that participants were sensitive to words’ bigram frequencies. Importantly, experiment 2 offers a conceptual replication of the results found in experiment 1. In sum, these results show that our design can capture participants’ sensitivity to mean bigram frequency, which is an effect typically studied in the context of orthographic processing.

### 3.2 Participants are sensitive to the co-occurrence of shapes in visual objects

Experiment 3 had exactly the same design as the preceding experiments, with the exception that the stimuli were now different from orthographic material. In fact, readers of Italian, who are used to a fully alphabetic script using Latin letters, were exposed to objects composed by a central Y-shaped body and a distinctive shape attached to each of three branches (similar to the stimuli used in [28], see Figure 2). While the overall objects play the role of words in experiments 1 and 2, the terminal shapes play a role analogous to characters.

We constructed the stimuli exactly as in experiment 1. We first selected 9 distinctive shapes, with which we constructed 6 standard objects. These objects were used as a base to construct the same two different types of deviants employed in the previous experiments. The first of such deviant objects was composed of pairs of shapes that were all present in the standard objects (high pair-frequency deviant). The second deviant object was instead constructed with pairs of shapes that were not present in any standard object (low pair-frequency deviant). Again similarly to experiment 1, different sets of images with the same statistical structure were created (two, in this case), and each participant was exposed to one of them. This had the goal of ruling out the possibility that the effects were driven by some idiosyncratic feature of a given set of shapes. An example stimuli set can be seen in Figure 2.

As in experiments 1 and 2, high pair-frequency deviants were harder to detect. The average *d’* for the high pair-frequency deviant was 0.65 [0.16, 1.15] and it was 1.58 [1.14, 2.03] for the low pair-frequency deviant. The difference in *d*’ between deviants was 0.93 [0.44, 1.43] (*t*_(38)_ = 3.81, *p* = 0.00025, *g* = 0.64 [0.27, 1.01]). In this experiment, the effect of interest was present in 74.36% [57.87%, 86.96%] of the participants (29 out of 39, one-sided binomial test: *p* = 0.0017; see Figure 3C).

These results show an effect akin to the sensitivity to bigram frequencies that participants displayed with pseudo-orthographic material in experiments 1 and 2. Therefore, such effect can also be observed when participants are presented with novel visual stimuli that are clearly non-orthographic but look instead as three-dimensional visual objects.

### 3.3 Participants are sensitive to the co-occurrence of low level visual features

On their own, the results of experiment 3 show that participants’ sensitivity to bigram frequencies can be observed outside the domain of orthographic processing. Yet, the stimuli were still similar to reading material insofar as higher-level units (the words in experiments 1 and 2, and the objects as wholes in experiment 3) were made up of lower-level parts (letters in experiments 1 and 2, and the terminal shapes in experiment 3) arranged in space. The stimuli used in experiment 4, instead, were circular sinusoidal gratings defined by combinations of low level visual features. These features (which played the role of characters in experiments 1 and 2, and of the terminal shapes in experiment 3) were spatial frequency, orientation and contrast.

In experiment 1, each of the 3 character positions defining a word could be occupied by 1 out of 3 possible characters. In the same way, each low level visual feature defining the sinusoidal gratings could take 3 different values. We used these values to construct stimuli with a statistical structure in all identical to that of experiments 1 and 3. For example, the high pair-frequency deviant in Figure 2 shares spatial frequency and contrast with the top left standard, shares orientation and contrast with the middle left standard, and shares spatial frequency and orientation with the bottom left standard. So, this deviant shares all its “bigrams” with standard stimuli. Instead, the low pair-frequency deviant in Figure 2 is defined by feature values present in the other gratings, but in a unique combination. A different shuffling of feature values was used for each participant.

In experiment 4, mean *d*’ was 1.01 [0.42, 1.60] for the high pair-frequency deviant and 2.22 [1.69, 2.74] for the low pair-frequency deviant. Once more, high pair-frequency deviants were harder to detect. The difference in *d*’ between deviants was 1.21 [0.61, 1.81] (*t*_(34)_ = 4.12, *p* = 0.00011, *g* = 0.74 [0.33, 1.14]). Also in this case, the majority of the participants (82.86% [66.35%, 93.44%], 29 out of 35. One-sided binomial test: *p* = 5.8e-05) showed an effect in the direction of the hypothesis (see Figure 3D).

These results show that a “bigram frequency” effect is even present when what defines stimuli are non-spatially-segregated low level visual features that constitute the building blocks of most visual perception.

A summary of participants’ hit and false alarm rates in all experiments can be found in Table S1, in the Supplemental Information section.

### 3.4 Participant’s sensitivity to feature co-occurrence is of equivalent magnitude, across stimuli types

After showing that participants were sensitive to the co-occurrence of visual features in stimuli as diverse as words, objects and simple gratings, we proceeded to compare the magnitude of this effect across experiments. We predicted that if these effects are the result of a domain-general visual mechanism, then they should have equivalent magnitudes regardless of the stimuli used in the experiments.

Because we were interested in evidence for the null (i.e., the difference between high and low pair-frequency detection does **not** differ across experiments), we performed a series of Bayesian independent-samples tests (JZS Bayes Factor [29–31]) comparing the difference between deviant types’ *d*’ across experiments.

All comparisons across experiments yielded BF_01_ values above 3 (see Table 1), indicating substantial evidence in favour of the null, compared to the alternative hypothesis. Therefore, regardless of the type of stimuli used, participants’ performance was biased by the co-occurrence of features to a similar extent. This suggests that in all cases the mechanism in play might be the same.

**Table 1:** Comparison of participants’ sensitivity to feature co-occurrence across experiments. All comparisons across experiments yielded BF_01_ values above 3, which implies substantial evidence in favour of the null compared to the alternative hypothesis. Note that the CI of all effect sizes include 0.

| Comparison across stimuli types | $BF_{01}$ | Hedges' $g$ |
| --- | --- | --- |
| 3 character words vs Objects | 4.21 | 0.16 [-0.36, 0.68] |
| 3 character words vs Sinusoidal gratings | 4.89 | -0.02 [-0.56, 0.51] |
| 6 character words vs Objects | 3.37 | 0.24 [-0.27, 0.76] |
| 6 character words vs Sinusoidal gratings | 4.86 | 0.05 [-0.47, 0.58] |
| 3 character words vs 6 character words | 4.31 | -0.09 [-0.68, 0.49] |
| Objects vs Sinusoidal gratings | 4.43 | -0.16 [-0.62, 0.29] |

## 4 Discussion

### 4.1 Letter co-occurrence and models of visual word identification

In experiment 1 and 2, we provide new evidence that when readers are confronted with novel, unfamiliar words, they code for the statistics of co-occurrence between letters. These data sit well with recent work using a similar approach [32], and strengthens the idea that readers capitalise on letter co-occurrence statistics to learn to visually identify novel words. Completely new to this work is the finding that sensitivity to bigram frequency is independent of word length (see 3.4). This is particularly notable, because as the task that the participants faced required the recognition of words, it could be expected that they would rely more heavily on letter statistics when word memorization was more difficult, that is, with longer words. The fact that an effect of equivalent magnitude emerges regardless of word length, suggests that bigram coding is not a strategic, task-specific effect; rather, it seems to be an intrinsic aspect of novel word coding.

Several models of visual word identification embody the idea that orthographic processing builds on letter co-occurrence statistics. Seminal work on Parallel Distributed Processing models explicitly stated that sublexical representations depend on the orthographic properties of words [33]. More recently, models were proposed that suggests a hierarchy where higher-order units are based on the statistics of co-occurrence between lower level units [e.g., 13]. Some of these models have been entirely realised computationally [34, 35], and have proven to account for a number of core experimental findings in visual word identification. Some of these models explicitly commit to the existence of bigram representations [13]; this is obviously in line with the results illustrated here, although we did not contrast different coding schemes (e.g., edit distance; spatial coding, [36]) and therefore we cannot speak in this respect.

### 4.2 Visual “word” identification of non-orthographic stimuli

While the models mentioned in the previous section can successfully account for a number of experimental findings in the context of reading, including the results of our experiments 1 and 2, they share a common limitation. By dealing only with orthographic stimuli, they do not to place reading within the broader context of visual perception. As a result, they provide an account of reading as domain-specific (either explicitly or implicitly). In sharp contrast, the results of our experiments 3 and 4 strongly suggest that at least some of the mechanisms at play during visual word identification are not specialized for orthographic processing.

The presence of an effect akin to sensitivity to bigrams’ frequencies in the case of stimuli radically different from orthographic material implies that the mechanism at play might be domain-general. This is in agreement with the proposal that cultural inventions such as writing recycle preexisting domain-general cognitive processes [2, 8]. Furthermore our results suggest that in the case of reading, the particular mechanism that is recycled is the visual system’s ability to extract statistical regularities in the co-occurrence of lower level units, and integrate them into higher level units.

Our intention is not to claim that every aspect of visual word identification is domain-general in nature. Certainly some effects found in the literature seem to reveal letters’ special status. For example, transposed-letter strings (e.g., NDTF for NTDF) are considerably more confusable than strings of non-letters symbols (e.g., %$&£ for %&$£; [37]); the accuracy profile in letter identification across different positions within strings is unique to letters (e.g., [38]); and attention is preferably deployed to the beginning of letter strings, but not to the beginning of strings of other symbols (e.g., [39]). But note that the existence of effects that are specific to orthographic material does not justify the conception of visual word identification as purely domain-specific. A successful model of reading should account for these letter specific effects, but still do it within the broader framework of visual processing.

### 4.3 Word learning in the bigger context of vision

Our results are in agreement with two major classes of theoretical frameworks in the field of visual neuroscience. The first asserts that visual object information is extracted along a largely feedforward hierarchy of processing stages, where tuning for progressively more complex visual features is built up incrementally [40]. That is, units in a given layer of the hierarchy integrate inputs from units of the previous layer, so as to gain selectivity for the combination of features the input neurons encode – for instance, inputs from simple edge detectors can be combined to produce tuning for more complex shapes, such as corners and curved boundaries.

These ideas have been instantiated in a number of neural network models, starting from Fukushima’s Neocognitron [41] and Riesenhubber’s and Poggio’s HMAX model [42], to arrive to modern deep convolutional neural networks [43]. These models not only have achieved extraordinary accuracies at classifying visual images, but have also been able to explain key trends in the tuning properties of ventral neurons in humans, monkeys and rats [44–49], as well as human and monkey performance in object recognition tasks [50–55]. Our findings are highly consistent with these models, since they show how distinct visual features (no matter whether characters, shapes, or low-level visual properties) are hierarchically integrated into progressively more complex combinations (e.g., bigrams or pairs of features) before being represented as full “objects” (e.g., whole words).

Interestingly, in our experiments such hierarchical feature integration took place spontaneously, via purely passive exposure to the statistics of the stimulus set. This resonates with another important theoretical principle that has been called into cause to explain why visual neurons develop certain kind of tuning properties and not others. This principle postulates that the tuning of sensory neurons is determined by adaptation to the statistics of the signals they need to encode [56–58]. Starting from the pioneering intuitions of Attneave and Barlow [59, 60], such efficient coding principle has been instantiated in a number of computational models that are able to learn key properties of the visual system through unsupervised exposure to the spatiotemporal regularities of the visual input [61–63].

Most notably, solid causal evidence has been gathered to show that visual neurons adaptively change their tuning depending on the statistics of the visual stimuli they have been passively exposed to, both during postnatal development [64–66] and adult life [67, 68]. At the perceptual level, human sensitivity to higher-order image statistics has been shown to closely match the informational content of such statistics in natural scenes [69, 70], thus suggesting a developmental adaptation of visual perception to the regularities of the visual world.

Moreover, passive exposure to altered object statistics has been shown to affect human performance in object recognition tasks [53, 71, 72], thus suggesting that unsupervised adaptation to visual input statistics continuously sculpt visual perception, even in adult life. Our findings add further behavioral evidence to this conclusion, by showing that enhanced sensitivity to specific pairs of visual features emerges as a result of their mere frequency of occurrence within a given stimulus set that is passively experienced by human observers.

### 4.4 Final conclusions

The results presented in this work suggest that a fundamental processing mechanism behind the learning of visual words also supports the learning of other combinations of visual objects/features. This implies that such mechanism is indeed of general-purpose, and confirms the view that reading builds on evolutionarily older cognitive structures [2, 8]. This mechanism would enable the statistical learning of regularities in the visual environment [e.g., 56, 58, 73, 74]. In this view, specialization for letter and letter clusters would emerge in skilled readers via the heavy exposure to written language that is characteristic of modern society [e.g., 75, 76]. To conclude, we hope this work will help to inspire models of reading that profit from the body of knowledge amassed in the broader field of visual neuroscience.

## 5 Acknowledgments

This research was supported by the ERC Starting Grant no. 679010-STATLEARN (DC) and by the ERC Consolidator Grant no. 616803-LEARN2SEE (DZ).

## 6 Author Contributions

Conceptualization: YV, EV, DZ & DC

Data curation: YV

Formal analysis: YV

Funding acquisition: DZ & DC

Investigation: YV & EV

Methodology: YV

Resources: YV & EV

Software: YV

Supervision: YV & DC

Visualization: YV

Writing original draft: YV & DC

Writing, review & editing: YV, EV, DZ & DC

## 8 STAR Methods

### 8.1 Resource Availability

#### 8.1.1 Lead contact

Further information and requests for resources and reagents should be directed to and will be fulfilled by the Lead Contact, Davide Crepaldi (, @CrepaldiDavide).

#### 8.1.2 Materials availability

The stimuli used in experiments 1, 2 and 3 can be found at the following Open Science Framework (OSF) repository: osf.io/3tyeu/

The stimuli used in experiment 4 were generated programmatically.

#### 8.1.3 Data and code availability

The data from all experiments, as well as the code used to perform the statistical analysis can be found at osf.io/3tyeu/.

### 8.2 Experimental Model and Subject Details

#### 8.2.1 Participants

All participants were self-reported right handed, Italian native speakers, and were recruited from the city of Trieste via on-line advertisement. They all had normal or corrected-to-normal vision and no language-related impairments. Participants provided informed consent and received a monetary compensation of 10 €. The experiment was approved by SISSA’s Ethical Committee.

Twenty-two participants (5 male and 17 female) took part in experiment 1 (mean age = 23.4, *σ* = 2.21 years); 25 participants (9 male, 16 female) took part in the experiment 2 (mean age = 25.04, *σ* = 3.1 years); 40 participants (12 male, 28 female) took part in experiment 3 (mean age = 24.22, *σ* = 2.57 years) and 36 participants (8 male, 28 female) took part in experiment 4 (mean age = 23.22, *σ* = 3.37 years).

### 8.3 Method Details

#### 8.3.1 Procedure and experimental design

The 4 experiments presented in this work followed the same procedure and used the same experimental design, differing only in the stimuli used. Participants sat in a sound-attenuated testing booth at around 70cm of a 27” computer monitor (BenQ XL2720Z). The experiment was programmed and run in MATLAB (2015b, MathWorks, Inc., Natick, MA, USA) using the Psychophysics Toolbox (v3) extensions [77, 78].

As described in the Results section, participants first completed a learning block where 200 standard stimuli were presented. During this block they were instructed to pay attention and try to learn the stimuli. After the learning block, participants completed 6 testing blocks, where standard stimuli were presented intermixed with deviant stimuli. On each of these blocks, participants were presented with 180 standard stimuli (90% of the trials; 15% each of the 6 standard tokens) and 20 deviant stimuli (10% percent of the trials; 5% each deviant condition).

Stimuli order of presentation was pseudorandom. Each test block started with 12 standard stimuli, after which deviants were presented allowing 6 to 12 standard stimuli in between each deviant presentation. Stimuli repetitions were only allowed after 2 other stimuli were presented (e.g., s2 s3 s1 s2).

Participants had a maximum of 2 seconds to classify each stimulus as either ‘Correct” (standard) or ‘Mistaken” (deviant) by pressing one of two keys on a keyboard. Key mapping was counterbalanced across participants. In case of timeout, the next trial started without any feedback. Overall, each participant was asked to classify 1080 instances of standard stimuli, and 60 instances of each deviant stimuli. Each block lasted on average 7 minutes and the entire experiment had an approximated duration of 50 minutes.

#### 8.3.2 Stimuli sets

The stimuli used in experiments 1 and 2 were words constructed using the Brussels Artificial Character Set (BACS [22]), whose characters have perimetric complexity, number of strokes, junctions and terminations matched to the English alphabet. We picked 23 out of the 26 available characters in BACS-2 with serifs. The 3 characters excluded were 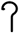 which resembles a 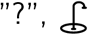, which is a vertical mirror flip image of 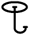, and 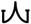 which resembles closely 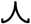.

The stimuli used in experiment 3 were not orthographic. Instead, we used images of 3D objects created using the software Blender (version 2.79b [79]).

Finally in experiment 4, the stimuli used were circular sinusoidal gratings defined by different values of three low level visual features. These features were spatial frequency (.4,.8 and 1.6 cycles per degree of visual angle), contrast (20%, 60% and 100%) and orientation (0, 45 and 90 degrees).

### 8.4 Quantification and Statistical Analysis

#### 8.4.1 Data and participants exclusion criteria

As we stated earlier, during the testing blocks participants had a time limit of 2 seconds to provide an answer. Trials in which participants did not provide an answer were excluded from the analyses, and participants with more than 20% of such trials for any stimulus category were excluded altogether.

In experiment 1, all participants provided enough trials in all conditions. Participants failed to provide an answer in 1.68% of the standard trials, 2.88% of the high pair-frequency deviant trials and 2.58% of the low pair-frequency deviant trials. Overall participants provided an answer in 98.21% of the trials. Two participants had to be excluded from the analysis in experiment 2. The remaining participants failed to provide an answer in 3.25% of the standard trials, 3.55% of the high pair-frequency deviant trials and 3.7% of the low pair-frequency deviant trials. Participants provided an answer in 96.71% of all trials. In experiment 3, one participant was excluded from the analysis. The rest of the participants failed to provide an answer in 2.05% of the standard trials, 2.74% of the high pair-frequency deviant trials and 2.65% of the low pair-frequency deviant trials. Of all trials, 97.88% were answered withing the time limit. Finally, one participant was excluded from the analysis in experiment 4. The remaining participants failed to provide an answer in 1.42% of the standard trials, 1.76% of high pair-frequency deviant trials and 2% of the low pair-frequency deviant trials. Participants provided an answer in 98.53% of all trials.

#### 8.4.2 Measure of performance

To better characterize the participants’ ability to detect deviant stimuli, we resorted to Signal Detection Theory and computed an independent sensitivity index (d-prime score or *d’)* for each deviant type and for each participant [25]. Participants’ responses were classified as “hit” (deviant stimuli classified as “mistaken”) or “false alarm” (standard stimuli classified as ‘‘mistaken”). Next, for each deviant type, *d*’ was calculated as

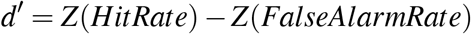

where Z(…) is the inverse of the cumulative standard normal distribution. This takes into consideration the overall bias towards a “correct” or a “mistaken” response. As this function does not output a finite value if either the hit rate or the false alarm rate are either 0 or 1, and considering the total amount of trials of each type, hit rate was capped between 1/60 and 59/60, and false alarm rate was capped between 1/1080 and 1079/1080.

#### 8.4.3 Statistical analysis

Statistical comparisons within each experiment were performed using paired-samples Student’s t-test when comparing *d*’ scores across deviants. Comparisons across experiments were performed using Bayesian independent samples tests (JZS Bayes Factor [29–31]). This test measures the relative evidence between the null and alternative hypotheses, allowing to assess evidence in favour of the null. Tests were performed using a Cauchy prior with scale value of *r* = 1. The code for this analysis was written by [80].

All effect sizes reported are Hedges’ *g* [26, 27], which is more precise than Cohen’s *d*, as it applies a correction for small sample sizes. Effect sizes were calculated using the Measures of Effect Size Toolbox [81]. All confidence intervals reported between square brackets are 95% CIs. In the case of Figure 3, as data is paired, the CI around the dot representing the mean performance at the group level is a within participants CI calculated using the normalization method proposed by Morey [82].

## 9 Supplemental Information

**Table SI:**
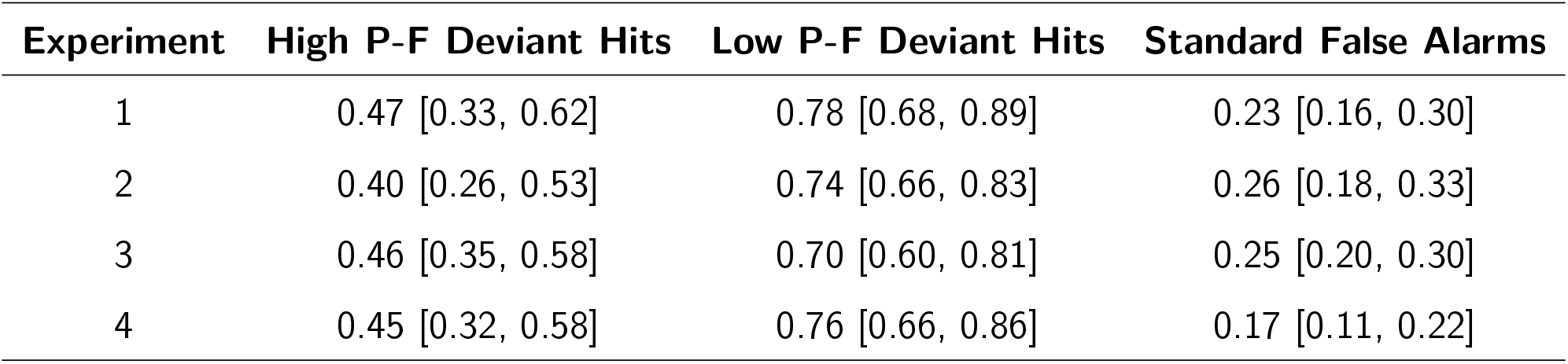
Hit rates and False Alarm rates in all experiments. As it can be seen, hit rates and false alarm rates were comparable across experiments.

## References

1. Woods, C., ed. (2010). Visible language (Chicago: Oriental Institute of the University of Chicago).

2. Dehaene, S. and Cohen, L. (2007). Cultural Recycling of Cortical Maps. Neuron 56, 384–398.

3. Dehaene, S., Pegado, F., Braga, L.W., Ventura, P., Nunes Filho, G., Jobert, A., Dehaene-Lambertz, G., Kolinsky, R., Morais, J., and Cohen, L. (2010). How learning to read changes the cortical networks for vision and language. Science 330, 1359–1364.

4. Ventura, P. (2014). Let’s face it: Reading acquisition, face and word processing. Frontiers in Psychology 5, 10–13.

5. Pegado, F., Bekinschtein, T., Chausson, N., Dehaene, S., Cohen, L., and Naccache, L. (2010). Probing the lifetimes of auditory novelty detection processes. Neuropsychologia 48, 3145–54.

6. Pegado, F., Comerlato, E., Ventura, F., Jobert, A., Nakamura, K., Buiatti, M., Ventura, P., Dehaene-Lambertz, G., Kolinsky, R., Morais, J., et al. (2014). Timing the impact of literacy on visual processing. Proceedings of the National Academy of Sciences of the United States of America 111, E5233–E5242.

7. Hervais-Adelman, A., Kumar, U., Mishra, R.K., Tripathi, V.N., Guleria, A., Singh, J.P., Eisner, F., and Huettig, F. (2019). Learning to read recycles visual cortical networks without destruction. Science Advances 5, eaax0262.

8. Dehaene, S. and Cohen, L. (2011). The unique role of the visual word form area in reading. Trends in Cognitive Sciences 15, 254–262.

9. Grainger, J., Dufau, S., Montant, M., Ziegler, J.C., and Fagot, J. (2012). Orthographic Processing in Baboons (Papio papio). Science 336, 245–248.

10. Scarf, D., Boy, K., Uber Reinert, A., Devine, J., Güntürkün, O., and Colombo, M. (2016). Orthographic processing in pigeons (Columba livia). Proceedings of the National Academy of Sciences 113, 201607870.

11. Rajalingham, R., Kar, K., Sanghavi, S., Dehaene, S., and DiCarlo, J.J. (2019). A potential cortical precursor of visual word form recognition in untrained monkeys. bioRxiv, DOI: 10.1101/739649.

12. Grainger, J. and Whitney, C. (2004). Does the huamn mnid raed wrods as a wlohe? Trends in Cognitive Sciences 8, 58–59.

13. Dehaene, S., Cohen, L., Sigman, M., and Vinckier, F. (2005). The neural code for written words: A proposal. Trends in Cognitive Sciences 9, 335–341.

14. Snell, J., van Leipsig, S., Grainger, J., and Meeter, M. (2018). OB1-reader: A model of word recognition and eye movements in text reading. Psychological Review 125, 969–984.

15. Chetail, F. (2015). Reconsidering the role of orthographic redundancy in visual word recognition. Frontiers in Psychology 6, 1–10.

16. Schmalz, X. and Mulatti, C. (2017). Busting a myth with the Bayes Factor. The Mental Lexicon 12, 263–282.

17. Binder, J.R., Medler, D.A., Westbury, C.F., Liebenthal, E., and Buchanan, L. (2006). Tuning of the human left fusiform gyrus to sublexical orthographic structure. NeuroImage 33, 739–748.

18. Vinckier, F., Dehaene, S., Jobert, A., Dubus, J.P., Sigman, M., and Cohen, L. (2007). Hierarchical Coding of Letter Strings in the Ventral Stream: Dissecting the Inner Organization of the Visual Word-Form System. Neuron 55, 143–156.

19. Lochy, A., Jacques, C., Maillard, L., Colnat-Coulbois, S., Rossion, B., and Jonas, J. (2018). Selective visual representation of letters and words in the left ventral occipito-temporal cortex with intracerebral recordings. Proceedings of the National Academy of Sciences of the United States of America 32, 1–10.

20. Maurer, U., Blau, V.C., Yoncheva, Y.N., and McCandliss, B.D. (2010). Development of visual expertise for reading: Rapid emergence of visual familiarity for an artificial script. Developmental Neuropsychology 35, 404–422.

21. Taylor, J.S., Plunkett, K., and Nation, K. (2011). The Influence of Consistency, Frequency, and Semantics on Learning to Read: An Artificial Orthography Paradigm. Journal of Experimental Psychology: Learning Memory and Cognition 37, 60–76.

22. Vidal, C., Content, A., and Chetail, F. (2017). BACS: The Brussels Artificial Character Sets for studies in cognitive psychology and neuroscience. Behavior Research Methods 46, 2093–2112.

23. Taylor, J.S.H., Davis, M.H., and Rastle, K. (2019). Mapping visual symbols onto spoken language along the ventral visual stream. Proceedings of the National Academy of Sciences of the United States of America, 201818575.

24. Endress, A.D. and Mehler, J. (2009). The surprising power of statistical learning: When fragment knowledge leads to false memories of unheard words. Journal of Memory and Language 60, 351–367.

25. Stanislaw, H. (1999). Calculation of signal detection theory measures. Behavior Research Methods, Instruments, & Computers 3, 37–149.

26. Hedges, L.V. (1981). Distribution Theory for Glass’s Estimator of Effect size and Related Estimators. Journal of Educational and Behavioral Statistics 6, 107–128.

27. Lakens, D. (2013). Calculating and reporting effect sizes to facilitate cumulative science: A practical primer for t-tests and ANOVAs. Frontiers in Psychology 4, 1–12.

28. Baker, C.I., Behrmann, M., and Olson, C.R. (2002). Impact of learning on representation of parts and wholes in monkey inferotemporal cortex. Nature Neuroscience 5, 1210–1216.

29. Rouder, J.N., Speckman, P.L., Sun, D., Morey, R.D., and Iverson, G. (2009). Bayesian t tests for accepting and rejecting the null hypothesis. Psychonomic Bulletin and Review 16, 225–237.

30. Jarosz, A.F. and Wiley, J. (2014). What Are the Odds? A Practical Guide to Computing and Reporting Bayes Factors. The Journal of Problem Solving 7, 2–9.

31. Leppink, J., O’Sullivan, P., and Winston, K. (2017). Evidence against vs. in favour of a null hypothesis. Perspectives on Medical Education 6, 115–118.

32. Chetail, F. (2017). What do we do with what we learn? Statistical learning of orthographic regularities impacts written word processing. Cognition 163, 103–120.

33. Pinker, S. and Prince, A. (1988). On language and connectionism: Analysis of a parallel distributed processing model of language acquisition. Cognition 28, 73–193.

34. Whitney, C. (2001). How the brain encodes the order of letters in a printed word: The seriol model and selective literature review. Psychonomic Bulletin and Review 8, 221–243.

35. Davis, C.J. (2010). SOLAR versus SERIOL revisited. European Journal of Cognitive Psychology 22, 695–724.

36. Davis, C.J. (2010). The spatial coding model of visual word identification. Psychological Review 117, 713–758.

37. Massol, S., Duñabeitia, J.A., Carreiras, M., and Grainger, J. (2013). Evidence for Letter-Specific Position Coding Mechanisms. PLoS ONE 8, 1–9.

38. Tydgat, I. and Grainger, J. (2009). Serial Position Effects in the Identification of Letters, Digits, and Symbols. Journal of Experimental Psychology: Human Perception and Performance 35, 480–498.

39. Scaltritti, M. and Balota, D.A. (2013). Are all letters really processed equally and in parallel? Further evidence of a robust first letter advantage. Acta Psychologica 144, 397–410.

40. DiCarlo, J.J., Zoccolan, D., and Rust, N.C. (2012). How does the brain solve visual object recognition? Neuron 73, 415–434.

41. Fukushima, K. (1980). Neocognitron: A self-organizing neural network model for a mechanism of pattern recognition unaffected by shift in position. Biological Cybernetics 36, 193–202.

42. Riesenhuber, M. and Poggio, T. (1999). Hierarchical models of object recognition in cortex. Nature Neuroscience 2, 1019–1025.

43. Lecun, Y., Bengio, Y., and Hinton, G. (2015). Deep learning. Nature 521, 436–444.

44. Cadieu, C., Kouh, M., Pasupathy, A., Connor, C.E., Riesenhuber, M., and Poggio, T. (2007). A model of V4 shape selectivity and invariance. Journal of Neurophysiology 98, 1733–1750.

45. Yamins, D.L., Hong, H., Cadieu, C.F., Solomon, E.A., Seibert, D., and DiCarlo, J.J. (2014). Performance-optimized hierarchical models predict neural responses in higher visual cortex. Proceedings of the National Academy of Sciences of the United States of America 111, 8619–8624.

46. Khaligh-Razavi, S.M. and Kriegeskorte, N. (2014). Deep Supervised, but Not Unsupervised, Models May Explain IT Cortical Representation. PLoS Computational Biology 10, e1003915.

47. Matteucci, G., Marotti, R.B., Riggi, M., Rosselli, F.B., and Zoccolan, D. (2019). Nonlinear processing of shape information in rat lateral extrastriate cortex. Journal of Neuroscience 39, 1649–1670.

48. Bashivan, P., Kar, K., and DiCarlo, J.J. (2019). Neural population control via deep image synthesis. Science 364, eaav9436.

49. Kietzmann, T.C., Spoerer, C.J., Sörensen, L.K., Cichy, R.M., Hauk, O., and Kriegeskorte, N. (2019). Recurrence is required to capture the representational dynamics of the human visual system. Proceedings of the National Academy of Sciences of the United States of America 116, 21854–21863.

50. Serre, T., Oliva, A., and Poggio, T. (2007). A feedforward architecture accounts for rapid categorization. Proceedings of the National Academy of Sciences of the United States of America 104, 6424–6429.

51. Rajalingham, R., Schmidt, K., and DiCarlo, J.J. (2015). Comparison of object recognition behavior in human and monkey. Journal of Neuroscience 35, 12127–12136.

52. Rajalingham, R., Issa, E.B., Bashivan, P., Kar, K., Schmidt, K., and DiCarlo, J.J. (2018). Large-scale, high-resolution comparison of the core visual object recognition behavior of humans, monkeys, and state-of-the-art deep artificial neural networks. Journal of Neuroscience 38, 7255–7269.

53. Cox, P.H. and Riesenhuber, M. (2015). There is a “U” in clutter: Evidence for robust sparse codes underlying clutter tolerance in human vision. Journal of Neuroscience 35, 14148–14159.

54. Kubilius, J., Bracci, S., and Op de Beeck, H.P. (2016). Deep Neural Networks as a Computational Model for Human Shape Sensitivity. PLoS Computational Biology 12, 1–26.

55. Kheradpisheh, S.R., Ghodrati, M., Ganjtabesh, M., and Masquelier, T. (2016). Deep Networks Can Resemble Human Feed-forward Vision in Invariant Object Recognition. Scientific Reports 6, 1–24.

56. Simoncelli, E.P. and Olshausen, B.A. (2001). Natural Image Statistics and Neural Representation. Annual Review of Neuroscience 24, 1193–1216.

57. Olshausen, B.A. and Field, D.J. (1997). Sparse Coding with an Overcomplete Basis Set: A Strategy Employed by V1? Coding V1 Gabor-wavelet Natural images. Vision Research 37, 3311–3325.

58. Olshausen, B.A. and Field, D.J. (2004). Sparse coding of sensory inputs. Current Opinion in Neurobiology 14, 481–487.

59. Attneave, F. (1954). Some informational aspects of visual perception. Psychological Review 61, 183–193.

60. Barlow, H.B. (1961). Possible Principles Underlying the Transformations of sensory messages. In Sensory Communication, W.A. Rosenblith, ed. (Cambridge MA: MIT Press), pp. 217–234.

61. Földiák, P. (1991). Learning Invariance from Transformation Sequences. Neural Computation 3, 194–200.

62. Olshausen, B.A. and Field, D.J. (1996). Emergence of simple-cell receptive field properties by learning a sparse code for natural images. Nature 381, 607–609.

63. Berkes, P. and Wiskott, L. (2005). Slow feature analysis yields a rich repertoire of complex cell properties. Journal of Vision 5, 579–602.

64. Hsu, A.S. and Dayan, P. (2007). An unsupervised learning model of neural plasticity: Orientation selectivity in goggle-reared kittens. Vision Research 47, 2868–2877.

65. Matteucci, G. and Zoccolan, D. (2020). Unsupervised experience with temporal continuity of the visual environment is causally involved in the development of V1 complex cells. Science Advances 6, eaba3742.

66. Arcaro, M.J., Schade, P.F., Vincent, J.L., Ponce, C.R., and Livingstone, M.S. (2017). Seeing faces is necessary for face-domain formation. Nature Neuroscience 20, 1404–1412.

67. Li, N. and DiCarlo, J.J. (2008). Unsupervised natural experience rapidly alters invariant object representation in visual cortex. Science 321, 1502–1507.

68. Li, N. and DiCarlo, J.J. (2010). Unsupervised natural visual experience rapidly reshapes size-invariant object representation in inferior temporal cortex. Neuron 67, 1062–1075.

69. Tkačik, G., Prentice, J.S., Victor, J.D., and Balasubramanian, V. (2010). Local statistics in natural scenes predict the saliency of synthetic textures. Proceedings of the National Academy of Sciences of the United States of America 107, 18149–18154.

70. Hermundstad, A.M., Briguglio, J.J., Conte, M.M., Victor, J.D., Balasubramanian, V., and Tkačik, G. (2014). Variance predicts salience in central sensory processing. eLife 3, 1–28.

71. Wallis, G. and Bülthoff, H.H. (2001). Effects of temporal association on recognition memory. Proceedings of the National Academy of Sciences of the United States of America 98, 4800–4804.

72. Perry, G., Rolls, E.T., and Stringer, S.M. (2006). Spatial vs temporal continuity in view invariant visual object recognition learning. Vision Research 46, 3994–4006.

73. Frost, R., Armstrong, B.C., and Christiansen, M.H. (2019). Statistical learning research: A critical review and possible new directions. Psychological Bulletin 145, 1128–1153.

74. Saffran, J.R., Aslin, R.N., and Newport, E.L. (1996). Statistical learning by 8-month-old infants. Science 274, 1926–1928.

75. Perea, M., Moret-Tatay, C., and Panadero, V. (2011). Suppression of mirror generalization for reversible letters: Evidence from masked priming. Journal of Memory and Language 65, 237–246.

76. Perea, M., Mallouh, R.A., García-Orza, J., and Carreiras, M. (2011). Masked priming effects are modulated by expertise in the script. Quarterly Journal of Experimental Psychology 64, 902–919.

77. Brainard, D. (1997). The Psychophysics Toolbox. Spatial Vision 10, 433–436.

78. Pelli, D. (1997). The VideoToolbox software for visual psychophysics: transforming numbers into movies. Spatial Vision 10, 437–442.

79. Blender Online Community (2017). Blender -a 3D modelling and rendering package.

80. Schwarzkopf, S. (2015). Bayes Factors Matlab functions.

81. Hentschke, H. and Stüttgen, M.C. (2011). Computation of measures of effect size for neuroscience data sets. European Journal of Neuroscience 34, 1887–1894.

82. Morey, R.D. (2008). Confidence Intervals from Normalized Data: A correction to Cousineau (2005). Tutorials in Quantitative Methods for Psychology 4, 61–64.

